# Temporo-nasally biased moving grating selectivity in mouse primary visual cortex

**DOI:** 10.1101/708644

**Authors:** Marie Tolkiehn, Simon R. Schultz

**Affiliations:** Centre for Neurotechnology and Department of Bioengineering, Imperial College London, SW7 2AZ, UK

**Keywords:** Mouse Primary Visual Cortex, Electrophysiology, Multi-Unit Activity, Salt-and-Pepper

## Abstract

Orientation tuning in mouse primary visual cortex (V1) has long been reported to have a random or “salt-and-pepper” organisation, lacking the structure found in cats and primates. Laminar in-vivo multi-electrode array recordings here reveal previously elusive structure in the representation of visual patterns in the mouse visual cortex, with temporo-nasally drifting gratings eliciting consistently highest neuronal responses across cortical layers and columns, whilst upward moving gratings reliably evoked the lowest activities. We suggest this bias in direction selectivity to be behaviourally relevant as objects moving into the visual field from the side or behind may pose a predatory threat to the mouse whereas upward moving objects do not. We found furthermore that direction preference and selectivity was affected by stimulus spatial frequency, and that spatial and directional tuning curves showed high signal correlations decreasing with distance between recording sites. In addition, we show that despite this bias in direction selectivity, it is possible to decode stimulus identity and that spatiotemporal features achieve higher accuracy in the decoding task whereas spike count or population counts are sufficient to decode spatial frequencies implying different encoding strategies.

**Significance statement:** We show that temporo-nasally drifting gratings (i.e. opposite the normal visual flow during forward movement) reliably elicit the highest neural activity in mouse primary visual cortex, whereas upward moving gratings reliably evoke the lowest responses. This encoding may be highly behaviourally relevant, as objects approaching from the periphery may pose a threat (e.g. predators), whereas upward moving objects do not. This is a result at odds with the belief that mouse primary visual cortex is randomly organised. Further to this biased representation, we show that direction tuning depends on the underlying spatial frequency and that tuning preference is spatially correlated both across layers and columns and decreases with cortical distance, providing evidence for structural organisation in mouse primary visual cortex.

## Introduction

Visual information processing in cortical circuits is still poorly understood. Electrophysiological studies in cat visual cortex area 17 revealed half a century ago that moving bars at different orientations evoked responses of varying strength (Hubel and Wiesel 1962). Hubel and Wiesel discovered orientation-selective neurons (Hubel and Wiesel 1962), and their organisation in orientation columns (Hubel and Wiesel 1974), consisting of neurons of the same or similar preferred orientation across multiple cortical layers. In addition to these orientation columns, orientation-selectivity was found to be organised laterally (Bonhoeffer and Grinvald 1991), where preferred orientation progressed in small increments, lateral to the cortical surface, covering the entire orientation field (Espinosa and Stryker 2012). These orientation maps are found in higher-level mammals such as cats, ferrets, or monkeys (Koulakov and Chklovskii 2002), but are apparently missing in rodents such as rats and mice.

Rat and mouse V1 do display orientation selectivity, with cells’ preferred orientations arranged randomly in what has been termed a “salt-and-pepper” organisation (Ohki, Chung, Ch’ng, Kara, and Reid 2005; Carrillo-Reid, Miller, Hamm, Jackson, and Yuste 2015; Chklovskii and Koulakov 2004; Kaschube 2014; Tan, Brown, Scholl, Mohanty, and Priebe 2011). Hansel (Hansel and van Vreeswijk 2012) argued random connectivity would be enough to create orientation selectivity in animals lacking topographic organisation, in contrast to the view that the generation of orientation selectivity required functional organisation (Corey and Scholl 2012). Recently, the question of orientation preference in mouse V1 was revisited, with some authors detecting more structure than hitherto reported (Kondo, Yoshida, and Ohki 2016; Ringach, Mineault, Tring, Olivas, Garcia-Junco-Clemente, and Trachtenberg 2016), with preferred orientations being found to show some degree of clustering both on cross-columnar and laminar scales. The median similarity in tuning preference was found to be significantly higher between neurons in close proximity (<100 *µ*m) than for neurons separated by 200 *µ*m (Ringach, Mineault, Tring, Olivas, Garcia-Junco-Clemente, and Trachtenberg 2016). Correspondingly, we hypothesise that orientation tuning preference and selectivity in mouse V1 is more correlated across layers and cortical columns than suggested by the salt-and-pepper tenet.

To test this, we analysed multi-unit activity (MUA, the aggregate spiking of a local population of cells, recorded by thresholding the signal on a single electrode site, but not spike-sorting), to examine both cross-laminar and cross-columnar arrangement of signals. Using multi-shank laminar electrodes, we recorded signals from electrode sites at different locations spaced 100 *µ*m apart in depth, and 200 *µ*m laterally, showing activity across layers and columns, effectively creating an electrophysiological “image” of the signals in the tissue, with the “pixel” size approximately corresponding to the spatial scale on which the MUA signal changes. Because of this fine-scale random structure (Ohki, Chung, Ch’ng, Kara, and Reid 2005), it might be expected that the MUA signal in mouse visual cortex contains little information about the spatial structure of a stimulus beyond retinotopy.

Here, we studied how visual information is processed in mouse V1. Analysis of the electrophysiological data revealed that low SF gratings moving from the temporal to nasal visual field reliably evoked the highest neural responses across layers and columns, indicating a strong bias in direction selectivity. In contrast, upward moving gratings consistently elicited the lowest activities across sites. Tuning functions demonstrated a high signal correlation across cortical layers and columns for both directions and SF. These correlations between tuning functions decreased with distance between sites indicating spatial organisation. We further found that maximum firing rate at each recording site was affected by the underlying SF of the drifting gratings. Despite these biased responses, it was possible to decode direction and SF from the activity patterns across channels with a high degree of confidence, emphasising that single units might not be required for decoding tasks in mouse visual cortex. Together, our data indicate that mouse primary visual cortex displays a higher spatial organisation than previously thought.

## Materials and Methods

### Surgical methods

All experiments were designed, implemented and performed in accordance with the Animals (scientific procedures) Act 1986 (UK) and the Home Office (UK) Animal Care Guidelines and approved by the Imperial College Animal Welfare and Ethical Review Body under Project License PPL 70/7355 and personal licences. Mice were group-housed and kept in a reversed 12 hour dark/light cycle and recordings were performed during the early dark phase.

Experiments were performed on 12 young adult female C57BL/6 mice with mean age 2.2 months (range 43 to 82 days). Animals were sedated intraperitoneally (i.p.) with chlorprothixene (0.5 mg/kg, Sigma-Aldrich, UK) and anaesthetised with isoflurane (2% for induction, 1 - 2% for surgery, 1% for electrophysiology in 1.2% O_2_, Harvard Apparatus, UK). To maintain clear airways and avoid tracheal secretions, 0.3 mg/kg (Animalcare, UK) atropine sulphate was injected subcutaneously; and 2 mg/kg dexamethasone (Organon, UK) was administered subcutaneously to prevent oedema. Depth of anaesthesia was monitored by testing the pedal-withdrawal reflex. Body temperature was maintained at 37.1 ±0.5°C. To prevent corneal dehydration whilst maintaining clear optical transmission, the right eye was kept moist with silicone oil (Sigma-Aldrich) applied continually (typically 2-3 times during recordings). The left eye was treated with eye ointment (Allergan Lacrilube) and covered with black tape to avoid confounding effects due to the binocular visual zone or inputs from the contralateral eye. Ear bars, a custom-built nose cone and an incisor adaptor kept the head in position. We made an incision over the skull and removed the skin to expose the skull between Bregma and Lambda, approximately an area of 1 cm^2^. A ground screw was situated over the contralateral cerebellum.

The craniotomy was targeted to a location −3.55 mm posterior to bregma and 2.5 mm lateral from the midline, on the Paxinos and Franklin mouse brain atlas (Paxinos and Franklin 2012). This was then rescaled to the skull of each mouse in our study. The distance between bregma and lambda was measured stereotactically, and the corrected target location calculated as *x*_*LOC*_ = −3.55 mm* z[mm]/4.2 mm and *y*_*LOC*_ = 2.5*z[mm]/4.2 mm, with *z* being the measured bregma-lambda distance for each mouse, and 4.2 mm being the literature (Paxinos and Franklin 2012) average distance between them (mean 4.03 mm, standard deviation 0.22 mm). The corrected target area was marked with a permanent marker. A small craniotomy (1 mm diameter) was drilled for the ground screw with a hand-held dental drill (Osada Success 40, 0.5 mm Meisinger drill bit), the dura removed with a 27G needle, and the ground screw-socket complex (Precision Technology Supplies, M1.0×2.0 Slot Cheese Machine Screw DIN84 A2 St/St and socket connector, MILL MAX, 851-43-050-10-001000 connector, sip socket) gently inserted into the craniotomy. An elastic ring, cut from a syringe tip or a PVC tube was glued onto the target area, and a craniotomy of approximately 3 mm in diameter made inside the well, and the dura retracted. A 4×8-shank translaminar linear Neuronexus silicon microelectrode array (A4×8-5mm-100-200-177) was lowered slowly into the brain to a depth of 1060 *µm*, at a speed of a few tens of *µm*/s, thus generally covering layers 4 to 6, and partially covering layer 2/3. Once the electrode reached the required depth, it was left to settle for 20-30 minutes before recording began.

### Visual stimuli

Visual stimuli were delivered via a Samsung SyncMaster 2233Z, 22” LCD monitor (Wang and Nikolić 2011), at a refresh rate of 60 Hz. The stimuli on the gamma-corrected monitor were displayed in 8-bit grey scale, with a mean screen luminance of 46.93 cd/m^2^. Sinusoidal drifting gratings were presented in pseudorandom order, at full contrast, for 20 repetitions of each of 6 different spatial frequencies in 8 directions. Gratings were presented for 1 second, with a 1 s inter-stimulus interval (Fig. 1 A). The temporal frequency was constant at 1.6 Hz. Each stimulus set was interleaved with 1 minute of a gray screen (at mean luminance), in order to estimate ongoing activity (Kenet, Bibitchkov, Tsodyks, Grinvald, and Arieli 2003; Niell and Stryker 2008; Jurjut, Georgieva, Busse, and Katzner 2017). The LCD display was placed 25cm away from the mouse’s right eye, covering approximately a 60°×75° region of the visual field. Stimuli were generated with FlyMouse, custom-developed visual presentation software (Tang, Jimenez, Chakraborty, and Schultz 2016) based on the FlyFly MATLAB Psychophysics Toolbox-based interface developed at Uppsala University (http://www.flyfly.se/about.html), and customized by S Ardila Jimenéz and M Tolkiehn. The spatial frequencies of the gratings presented were [0.01, 0.02, 0.04, 0.08, 0.16, 0.32] cycles per degree (cpd), and directions were presented in 45 degree increments around the circle.

**Figure 1:**
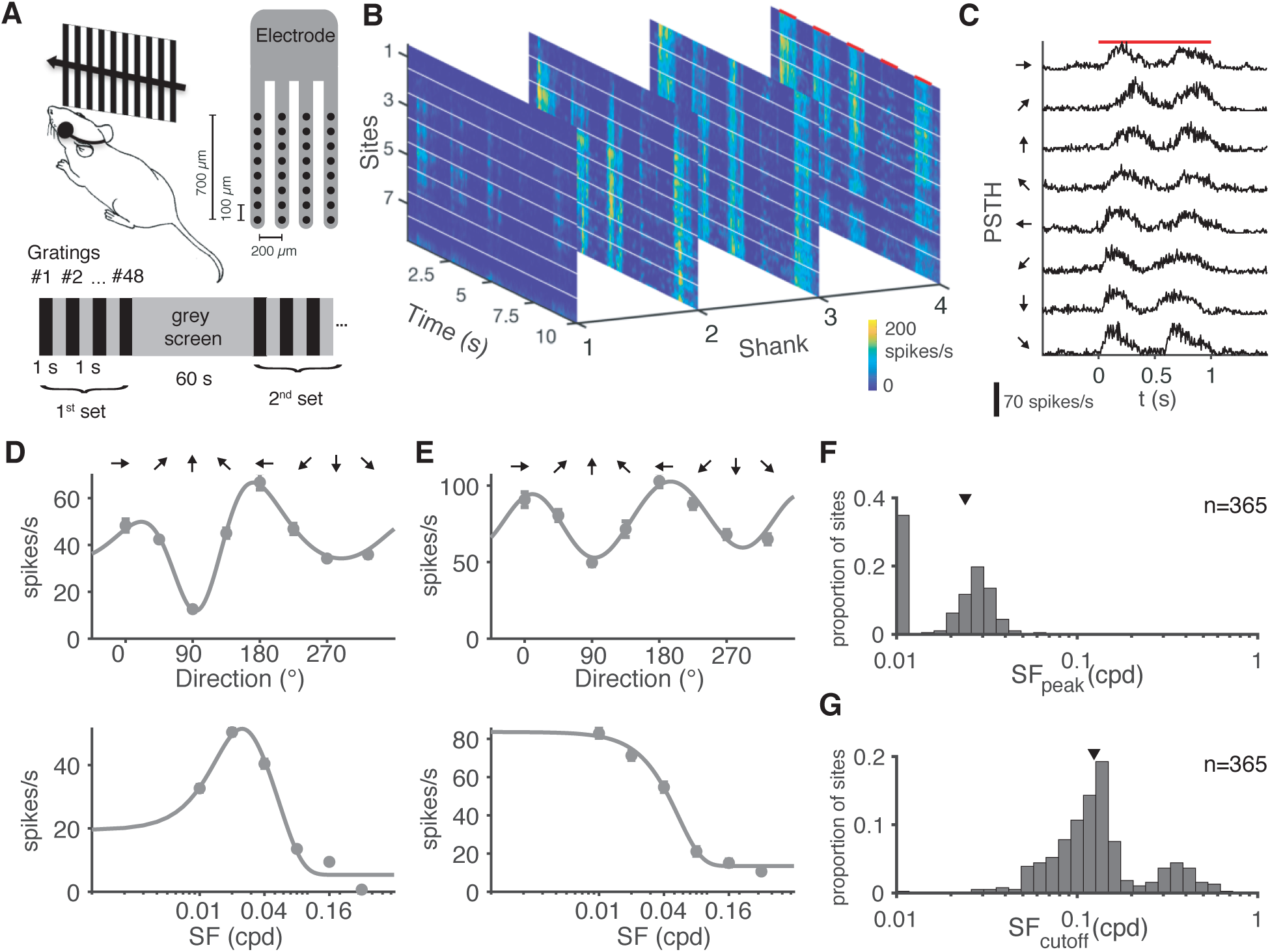
Grating stimuli strongly modulate activity across sites. (A) Top left: Moving gratings are presented on a monitor covering 60°×75° of the visual field of the right eye; in-vivo electrophysiological recordings were made from left hemisphere V1. Top right: cartoon of probe geometry. Bottom: Stimulus presentation (black) with interleaved grey screen at 1 s *on* and 1 s *off* time, in pseudorandom order, interleaved with 60 s grey screen. (B) Example electrophysiological “image”: colour-coded multi-unit firing rate for each site location, with 20 repetitions vertically stacked. 10 s of 5 different moving gratings (repetitions aligned) is shown. Red bars indicate stimulation, white lines separate recording sites. (C) Example site PSTH for 8 directions at 0.01 cpd, averaged over 20 repetitions and smoothed with a Hann window over 25 ms. Red bar indicates stimulation period. (D, E) Top: Von-Mises fits to baseline-subtracted data with directional peaks around 180° (left moving grating) for 2 example sites from 2 mice. Mean responses for directions were averaged across the two lowest SFs. Bottom: SF tuning for the same electrode sites. Note logarithmic x-axes. Responses to each SF were pooled over directions. Note different y-axis scales. Error bars denote s.e.m. (F) Distribution of peak SF. Triangle indicates median (0.023 cpd) over n=365 sites. (G) Distribution of −3 dB cut-off frequencies, with mode at 0.13 cpd. Triangle indicates median (0.12 cpd) over n=365 sites. Histograms in (E,F) use 30 bins logarithmically spaced between 0.01 and 1 cpd.

### Signal processing and data analysis

Signals were acquired with a Ripple Grapevine (Scout Processor), amplified with a single-reference amplifier with on-board filtering and digitization at 16 bit resolution and 0.2 *µ* V/bit (Grapevine Nano front end), using the Trellis software package. Broad-band signals sampled at 30 kHz were recorded, and filtered between 0.3 Hz and 7.5 kHz (3rd order Butterworth filter). The screen output signal and stimulus presentation triggers were synchronised with a custom-built photo-diode circuit board and photo-sensor (LCM555CN), attached to the bottom left corner of the monitor. A small rectangular field flashing at stimulus on- and offset was used as a synchronization pulse.

We high-pass filtered the electrophysiological data, and thresholded it at 4 standard deviations, to obtain the signals. Direction tuning of sites was evaluated using the sum of two modified von Mises functions (Gao, DeAngelis, and Burkhalter 2010; Swindale 1998; Gatto and Jammalamadaka 2007). SF tuning curves were fit with a Difference of Gaussiants (DoG) function (Grubb and Thompson 2003; So and Shapley 1981; Rodieck 1965). From these fits we calculated each site’s peak and cut-off spatial frequency, as well as preferred direction. Goodness of fit was estimated with the coefficient of determination, *R*^2^ = 1 − *sse*/*sstotal*, with *sse* denoting the sum of squared errors, and *sstotal* = (*n* − 1)*var*(*x*) the total variation. Sites with *R*^2^ <= 0.9 for both SF and directional fit were discarded from further tuning analysis, which were 14 out of 384 sites at 0.01 cpd (cf. Results).

For population tuning curve calculations, the site multi-unit firing rates (responses) were normalised across directions and SFs to fall between 0 and 1 for each repetition, enabling us to compare across channels while accounting for slow temporal changes in excitability and different site firing rates. Tuning curves were estimated by fitting the trial-averaged responses for each direction or SF. We then compared the fitted tuning functions across sites by calculating the pairwise Pearson correlation coefficient (*r*_*signal*_), as well as the noise correlation (*r*_*noise*_), estimated as the Pearson correlation of deviations of each trial from the mean response for that direction. Direction and orientation selectivity were calculated using the Direction Selectivity Index (DSI) and Orientation Selectivity Index (OSI) as described in (Mazurek, Kager, and Van Hooser 2014).

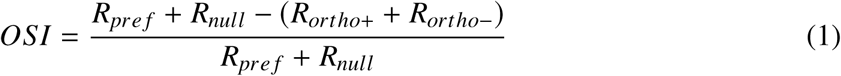

with *R*_*pref*_ as the preferred direction, *R*_*null*_ the opposite direction, and *R*_*ortho*±_ denoting the orthogonal directions. DSI was defined as:

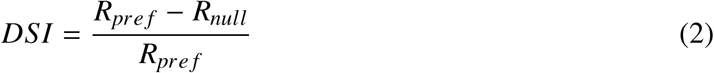

As DSI and OSI are known to be positively biased (Mazurek, Kager, and Van Hooser 2014), we also calculated measures related to circular variance *L*_*osi*_ and *L*_*dir*_ proposed by Mazurek et al.(Mazurek, Kager, and Van Hooser 2014):

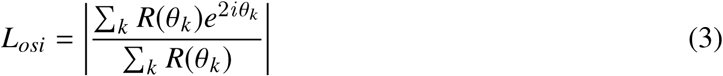

where circular variance is *CirVar* = 1 − *L*_*OSI*_. Correspondingly, a quantity related to circular variance in direction space via *DirCircVar* = 1 − *L*_*dir*_, scaled to fall between 0 and 1 as in Mazurek et al.:

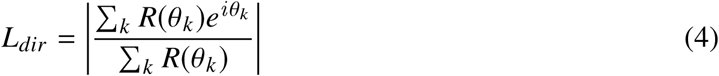

To investigate how stimulus information is encoded in the neural signal, and in particular, which features tell us more about certain aspects of the stimuli, three neural features were evaluated. As a basis, spikes of all sites were binned at 5 ms, binarised if multiple spikes occurred in one bin, creating a spatiotemporal matrix **M** of size 32×200 for each 1 s trial. Then, we used the following features: a) vectorising **M** to create spatiotemporal MUA (STMUA), a 6400×1 vector, b) spike count, where **M** was summed over the 1 second stimulus duration for each site (creating a 32×1 vector), and c) population rate (**M** was summed across sites, creating a 200×1 vector). STMUA allowed us to use all the spatiotemporal features in the data, spike count sums over the time course, greatly reducing the feature space to the number of channels, and summing over all channels provided us with the population rate, which retains the temporal information, yet loses information about site-specific contributions. We performed multinomial classification of the grating stimuli, using a Naive Bayes (NB) classifier to predict stimulus labels, implemented with MATLAB’s *fitcnb* function with a multinomial distribution input parameter in this task. The decoding task comprised two parts for each feature type: *A)* Decoding the SF (1 of 6), and *B)* Decoding the direction (1 of 8). 2-fold cross-validation was used. Training and testing data consisted of random uniformly sampled partitions of 50% training and 50% for testing, using MATLAB’s *datasample* function without replacement. All stimuli occurred with equal probability. For the SF decoding task, all training and testing data was taken at the same direction, 180°. For direction decoding, training and testing data was taken at 0.02 cpd. This was to ensure the stimuli were in a detectable range. Classification accuracy was evaluated against chance level, validated with 2-fold cross-validation and bootstrapped over 100 repetitions, with random permutations of the partitions, and across all mice. To do this, samples were drawn from the same random repetition set, to ensure training and testing samples were taken in close temporal coherence. Chance classification rates were tested with permutation tests (shuffling labels 1000 times).

### Data and code availability

Data and code of this study are available from the corresponding author upon reasonable request. Some of the data reported here has been previously described in (Tolkiehn and Schultz 2015). The current paper incorporates additional data and expands upon the analysis of the data.

## Results

We recorded multi-unit activity from the left primary visual cortex of 12 isoflurane-anaesthetised mice. Using 4-shank silicon microelectrodes with 8 linearly arranged recording sites per shank at 100 *µm* site and 200 *µm* shank spacing, activity was recorded from sites spanning the cortical laminar depth. Visual stimuli consisting of monocular, full-field drifting gratings were presented to the right eye (Fig. 1 (A), and Methods). Visual response properties including directional or spatial tuning were characterised and compared among sites in order to investigate visual information processing across cortical layers.

### Activity in V1 is highly modulated by drifting gratings

Multi-unit sites were strongly driven by the moving gratings, as illustrated in Fig. 1 (B) and in a post-stimulus time histogram (PSTH) in (C). From Fig. 1 (B) it is evident that the stimuli elicited responses of varying strength across site and shank locations (e.g. on shank 4, the 4th stimulus elicited weak activity and stronger in the 5th), with robust response magnitudes for each site across repetitions. We rejected sites from further analyses if the *R*^2^ of the tuning function fit fell below 0.9. The number of sites rejected in this manner was dependent upon the spatial frequency presented. Thus, at 0.01 cpd we used 370 of 384 sites, 371 at 0.02 cpd, 367 at 0.04 cpd, 354 at 0.08 cpd, 356 at 0.16 cpd, and 334 at 0.32 cpd. When pooling over all 6 SFs, we kept 344 sites, and at the two lowest SFs 369 sites. SF tuning was pooled over all directions, and we rejected 19 sites whose *R*^2^ fell below 0.9.

Fitting direction tuning curves (an additive pair of von Mises functions) to the baseline-subtracted, trial-averaged multi-unit responses of individual electrode sites revealed strong response modulation over multiple channels and shanks. Two examples of strongly tuned sites of different animals is presented in Fig. 1 (D, E, top), which show direction tuning curves averaged over spatial frequencies of 0.01-0.02 cpd (i.e. using spatial frequencies that should optimally evoke responses (Niell and Stryker 2008)). These examples were typical of the majority of sites across animals, which exhibited tuning functions similar in shape to those presented in Fig. 1 (D-E, top). This modulation was present at different average firing rates. In particular, the tuning functions of the examples reveal a global minimum at approximately 90°, and a maximum around 180°. It is also evident that the direction 180° appears to drive the activity slightly more than the collinear direction of 0°. Overall, the direction preferences of multi-unit sites tended to occur consistently at around 180°, with a median preferred direction of 182.2° (over n=370 of 384 sites) at SF 0.01 cpd.

Similar to direction tuning, the multi-unit activity was also modulated by the spatial frequency of the stimulus. In particular, the three lowest SFs (0.01 to 0.04 cpd) elicited high responses, whereas the three highest SFs induced lower activities. Difference-of-Gaussian (DoG) fits were estimated using the baseline-subtracted, trial-averaged responses (average across all presentations at the same SF). Individual sites were observed to have either bandpass (Fig. 1 (D, bottom)) or low-pass (E, bottom) properties, with low-pass defined by a monotone decreasing shape, and bandpass by a rise-fall shape in the amplitude-frequency plot. The overall mean preferred SF amounted to 0.02 cpd (s.e.m. = 0.0009 cpd, n=365).

The distribution of peak SF was bimodal, with a peak at the lowest SF (133 sites). Approximately two thirds of the sites (n=232) revealed bandpass properties indicated by a drop in response at the lowest SF and a secondary peak at 0.028 cpd, c.f. Fig. 1 (F). Most sites revealed a preferred SF of around 0.02-0.03 cpd. Variation was low and we only rarely observed a peak SF response exceeding 0.04 cpd (Fig. 1 (F)). In most sites, activity decreased drastically with SFs exceeding 0.04 cpd (Fig. 1 (F)), with a median −3-dB cut-off frequency of 0.12 cpd (Fig. 1 (G).

### Direction tuning in left V1 is biased towards leftward moving gratings and depends on spatial frequency

In the examples shown in Fig. 1 (D, E), the peaks and troughs of the direction tuning functions were closely aligned. To investigate if this was indicative of an overall bias in directional preference, we quantified which directions elicited the highest and lowest mean responses for each site across animals. If there was no underlying directional preference and a balanced representation, then the directions evoking maximum responses should average out over many samples and approximate a uniform distribution on the circle. Since average activity decreased substantially at high SFs (Fig. 1 (D, E bottom)), we investigated these metrics for each SF individually to compare mean responses at each SF and found that the distributions of mean response maxima and minima were distinctly affected by SF (Fig. 2 A, B).

**Figure 2:**
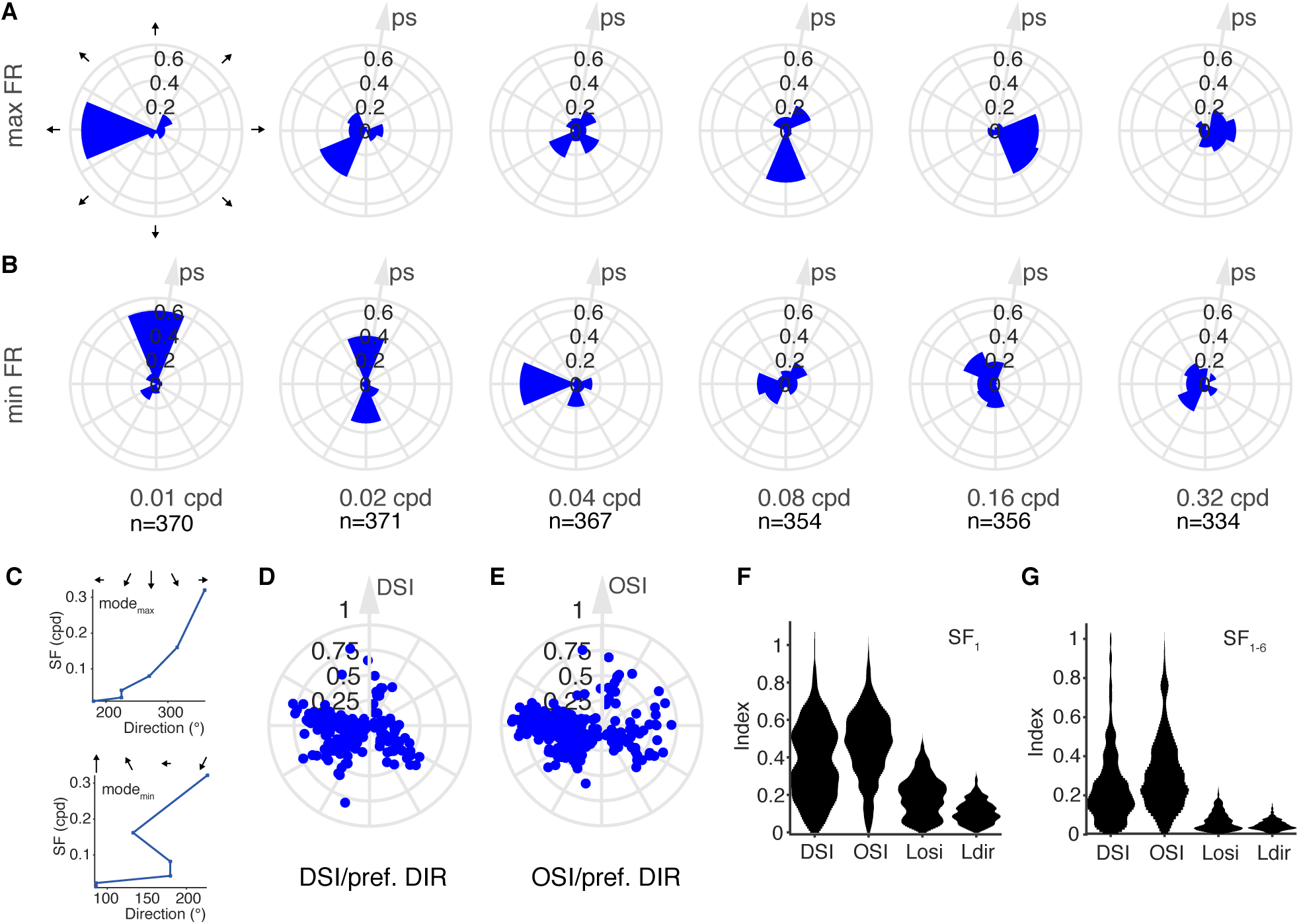
Maximum and minimum responses, and selectivity depend on spatial frequency. (A) Polar histograms of directions eliciting maximum responses of all multi-unit sites at each SF, with SFs in increasing order, left to right. Radial axis: proportion of sites (ps). (B) Polar histograms of directions eliciting minimum responses at each SF. (C) Top: Modes of maximum responses shown in (A). Bottom: Modes of the minimum responses shown in (B). (D) Polar plot of preferred direction (from peak tuning fit) against Direction Selectivity Index (DSI), where each dot represents the properties of one electrode site, pooled over 0.01-0.02 cpd. (E) Preferred direction against Orientation Selectivity Index (OSI). (F) Selectivity indices (DSI, OSI, *L*_*osi*_ and *L*_*dir*_) obtained from the lowest SF (0.01 cpd) alone. (G) Selectivity indices obtained by pooling all 6 SFs.

The distributions of directions evoking maximum and minimum were not uniformly distributed (Rayleigh test for non-uniformity of circular data, max: p = 5.7e-303, Z = 368.74, min: p = 1.6e-301, Z = 368.54, *α* = 0.05, n=369), but instead showed a high preference to leftward moving gratings (180°, corresponding to a movement from temporal to nasal visual field), which evoked maximum responses in the majority, 61.1%, of sites (Fig. 2 (A)). Upward moving gratings (90°) most frequently (59.7%) evoked minimal responses for low SFs (Fig. 2 (B)), as was also visible in the tuning functions of Fig. 1 (D, E top). The polarity of the proportion of sites decreased at higher SFs, with the effect of more uniform and less uniquely peaked polar histograms. Whilst for low SFs 40-70% of the sites grouped at similar (or collinear) directions, higher SFs appeared more evenly distributed with several directions accumulating levels around 15-20%.

The modes of the distributions revealed a distinct relationship between the directions that most frequently elicited the maximum response against each SF (Fig. 2 (C, top)), moving from 180° over 270° to 0° (however at decreasing polarity, cf. Fig. 2 (A)). The relationship between minimum response modes and SF was not as smooth as for the maximum response distributions, Fig. 2 (C, bottom), which was also visible in Fig. 2 (B). Together, the results highlight a strong direction selectivity and that particularly at 0.01 cpd, the directions eliciting maximum and minimum responses are highly biased across sites and animals.

This directional bias towards leftward moving gratings was also found in the preferred directions obtained by taking the peak of the von-Mises function fits, illustrated in Fig. 3 (D-E). Plotting the preferred directions against Direction Selectivity Index (DSI) and Orientation Selectivity Index (OSI) further illustrated this bias, showing it occurred in sites ranging from low to high direction (D) and orientation (E) selectivity. Direction selectivity was further explored with the more conservative stimulus selectivity estimates *L*_*osi*_ and *L*_*dir*_ (Fig. 2 (F-G)). The selectivity indices were also subject to the choice of SF: Estimating the selectivity indices at 0.01 cpd only as presented in (F) resulted in high values, at a median OSI 0.49, DSI 0.34, *L*_*osi*_ 0.20 and *L*_*dir*_ 0.10 (distributions over n = 370 sites). DSI and OSI distribution values ranged from 0.2 to 0.6, with DSI occupying lower indices than OSI, demonstrating a pronounced direction and orientation selectivity of the sites. The more conservative selectivity measures *L*_*osi*_ and *L*_*dir*_ occupied a much smaller range between 0 and 0.2, with direction selectivity centred around yet smaller values, concordant with OSI and DSI. 124 of 370 sites showed significant direction-selectivity (*α* = 0.01), estimated for *L*_*dir*_, as determined by a permutation test by reshuffling labels 1000 times. 277 of 370 sites were significantly orientation-selective on *L*_*osi*_ (permutation test, 1000 random sets, p < 0.01). Median null distributions for *L*_*dir*_ and *L*_*osi*_ were both 0.05.

**Figure 3:**
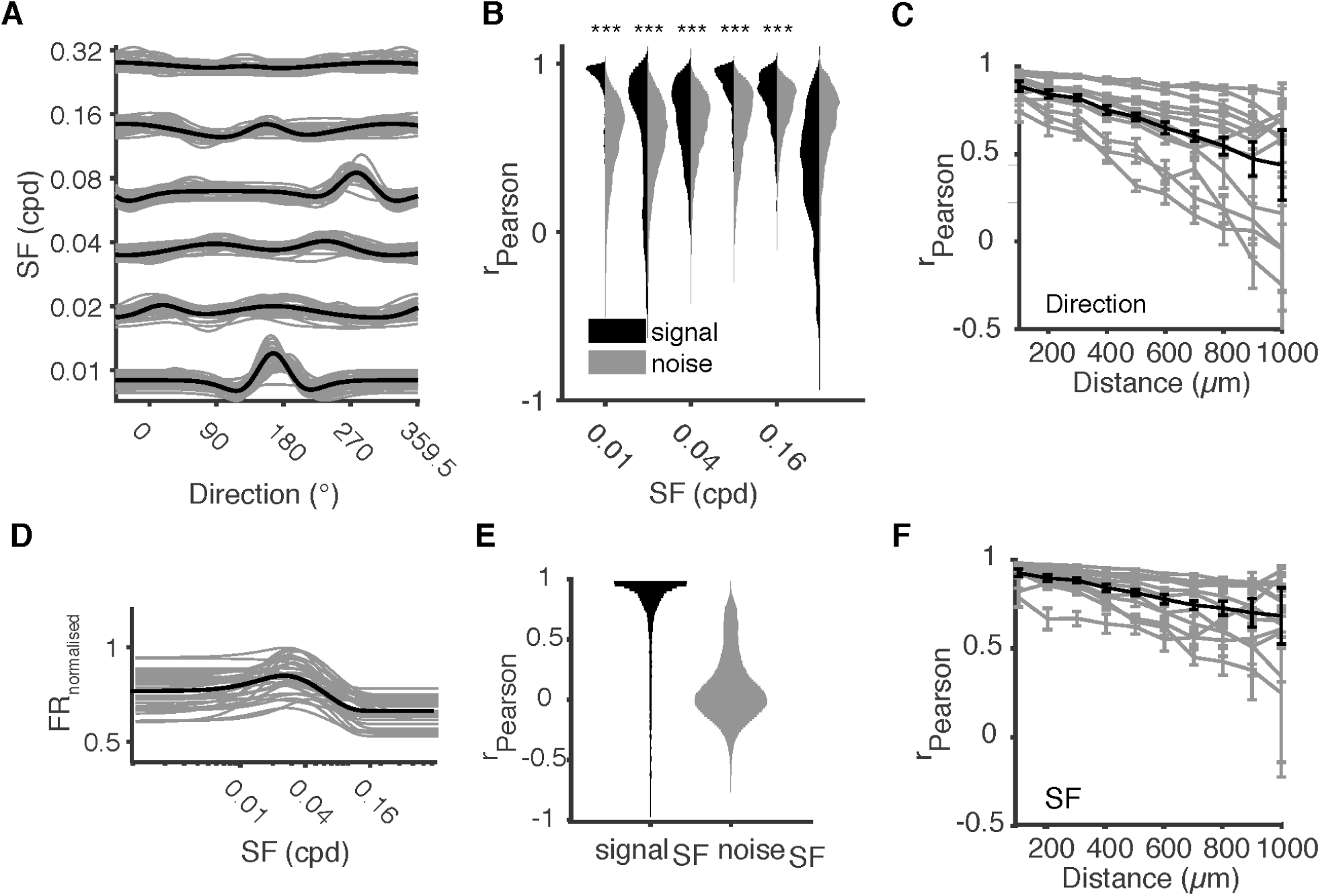
Direction tuning depends on underlying SF and is highly correlated between sites, even across shanks. (A) shows normalised tuning functions of all sites (grey, thin lines) for an example mouse, bottom to top: increasing SFs. Thick black line indicates fit on the mean over sites. (B) Pairwise signal (black) and noise (grey) correlation coefficients between directional tuning functions of all sites selected for analysis (*R*^2^ > 0.9, 31-32 sites) for the example mouse of (A). *** indicates p<0.001, one-sided Mann-Whitney-U. (C) Mean signal correlation between direction tuning functions across sites as a function of distance between sites, determined from probe geometry, discretised into 100 *µm* intervals. Each grey line represents one mouse, black line mean across mice; spatial frequency 0.01 cpd, error bars s.e.m. (D) Example mouse SF tuning function fits of normalised responses. Black line indicates tuning fit of the mean across normalised sites. (E) Pearson signal correlations between the fitted SF tuning curves at each electrode site (black), shown together with Pearson noise correlation (grey) in all mice (n=344 sites). (F) Mean signal correlation between SF tuning functions, estimated over the average across directions, as a function of distance between sites.

Estimating selectivity indices across all SFs resulted in a median DSI of 0.19 and a median OSI of 0.26; a median *L*_*osi*_ and *L*_*dir*_ of 0.06 and 0.04, respectively (estimated over n = 344 sites), as shown in Fig. 2 (G). 106 of 344 sites were significantly direction-selective, and 178 of 344 sites were significantly orientation-selective (permutation test, 1000 random shuffles, *α* = 0.01, p < 0.01, median selectivities of null distributions both 0.02).

### Tuning in V1 is highly correlated between sites

Direction tuning in V1 appeared to be distinctly correlated across sites, both longitudinally and laterally. The assumption of uniformly salt-and-pepper organised orientation maps requires the tuning functions to cover the whole direction space. Yet, peak responses appeared much more stereotyped and to be affected by the SF. Thus, we estimated the tuning functions of all sites after normalising their FRs and compared their shapes between sites and at different SFs. At 0.01 cpd, the peaks of the tuning functions seemed to align approximately with the example curves from Fig. 1 (D, E) across sites, while tuning function shapes at higher SFs differed (3 (A)).

We quantified the similarity of pairs of individual sites’ tuning functions within mice by calculating the (Pearson) signal and noise correlation coefficients. Across all mice, signal correlations between individual (within-mouse) tuning curves was high at low SF, with few negative correlations indicating opposing tuning curves (Fig. 3 (B)). Noise correlations were generally significantly lower (one-sided Mann-Whitney U test, Fig. 3 (B)). Comparison of Figs. 3 (A) and (B) revealed that in Fig. 3 (A), tuning appeared diverse on the SFs, with a very high signal correlation at the lowest SF tightly located at a correlation coefficient of 1, and a large spread over many values at high SFs, while noise correlations appeared similar across SFs.

Given these high signal correlations between sites and our probe geometry, we next investigated if this correlation depended on site distance. When comparing individual neurons, the salt-and-pepper notion stipulates that there should not be a relationship between tuning similarity and distance, as each neuron, regardless of its location should display a random direction preference and no clustering. Assuming MUs pool over a large number of neurons, each electrode site would be expected to show identical responses regardless of the stimulus bar the SF cut-off. Thus, in both cases, correlation should not be affected by distance. However, our data shows the average Pearson (signal) correlation between direction tuning functions decreased with distance between sites (Fig. 3 (C)), where distance was inferred by probe geometry and discretised into 100 *µ*m bins.

These findings were consonant with the normalised DoG tuning functions for SFs, demonstrating high similarity over sites and shanks (Fig. 3 (D)). Correlation coefficients for SF tunings appeared all high with a centre of mass near a correlation coefficient of 1, as evident from Fig. 3 (E). Only a few strongly opposing negative values were visible, and noise correlations appeared generally centred around zero. Signal correlations were significantly higher than noise correlations (median correlation signal 0.92, noise 0.07, two-sided Mann-Whitney U, *α* = 0.05, Z = 112.78, n_*signal*_ = 5952 (12 mice at each 32 pairwise 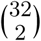 combinations), n_*noise*_ = 35712, p < 10^−100^), as evident from the distributions in Fig. 3 (F), demonstrating the difference between noise and driven activity.

This was also confirmed by the average Pearson (signal) correlations between spatial tuning functions, Fig. 3 (F). Yet, the spatial tuning similarity decreased more slowly than the tuning similarity of direction tuning with distance.

### High decoding success in spatial frequency and direction decoding

Given the high correlation between both SF and direction tuning and the bias towards leftward forward moving gratings particularly for low SFs, we investigated if and how well different features of multi-unit activity could be used to decode the stimulus features.

To do that, we devised two multivariate decoding tasks using Naive Bayes decoder: a) SF decoding and b) direction decoding. Both decoding tasks were designed in balanced sets: The decoder was trained on 50% of the data (10 trials for each decoding target), and tested on the remaining 50% (10 trials for each decoding target). Decoding was done per individual mouse, 2-fold cross-validated and bootstrapped 100 times for random subsets of the data. Each of the 100 bootstrapping steps in the SF decoding task involved 60 samples (10 samples at 6 SF), and decoding required to correctly identify one in 6 SFs (chance level 16.7%), each at the same direction of 180°. Equivalently, in the direction decoding task, for each bootstrap set 80 samples (10 samples for all 8 directions) were classified at 0.02 cpd.

Using either spatiotemporal multi-unit activity (STMUA), spike count (SC) or population rates (Pop), we achieved high classification rates of around 50% for SF decoding. Fig. 4 (A) shows the mouse-averaged confusion matrix for decoding SF from spike responses (STMUA), where circle size and colour correspond to percentage correctly classified samples. Low SFs scored a high decoding accuracy, while higher SFs tended to be underestimated, indicated by higher misclassifications below the identity line (max 87% for 0.01 cpd). Panel (B) of Fig. 4 reveals the mouse-averaged confusion matrix for direction decoding achieving an evenly high classification performance across all directions (max 81% at 225°).

**Figure 4:**
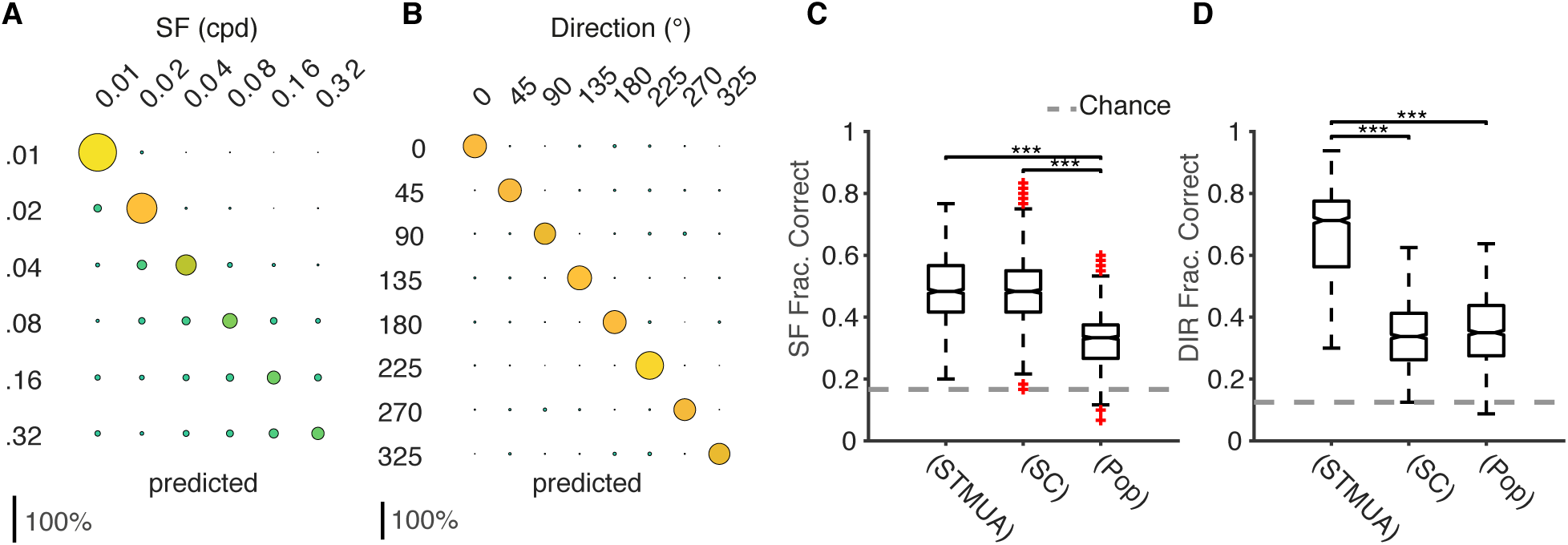
SF and direction can be accurately decoded from multi-unit neural signals. (A, B) Mouse-averaged confusion matrices (predictions x-axis, and actual labels y-axis) for decoding (A) SF and (B) direction from the STMUA. Circle radius and colour correspond to correctly classified samples each of size 32 channels and 200 bins of 5 ms. (C) Correct classification rates for all mice for SF-decoding, from the binary spike train vector (STMUA), the vector of spike counts of all channels (SC) and the population rate (Pop). 2-fold cross-validation used for each mouse, with 100x bootstrapping. Dashed grey line indicates chance at 16.7%. (D) As in (C) but for direction decoding, chance at 12.5%. ** indicates p<0.01 and *** p<0.001, one-sided sign test, *α*=0.05, Bonferroni corrected.

In the SF decoding task, the bootstrapped, 2-fold cross-validated decoding performances revealed a median performance of 48.1% for STMUA, the spike count features achieved 50.7% and population rate features 31.6% (Fig. 4 (C)). In the box plots of (C-D), the central mark represents the median, the bottom and top edges the 25th and 75th percentiles. The top and bottom whiskers indicate the extreme data points which are not considered outliers, while the outliers are indicated individually in red crosses. Direction decoding attained a median of 73.3% (STMUA), spike count features 34.5%, and population rate 37.1% (against a chance level of 12.5%), as presented in Fig. 4 (D).

All results showed statistically significant classification performance, as revealed by permutation tests (1000 random permutations with shuffled labels, median across mice all p<0.001). Median of shuffled tests was 12.5% for direction decoding, and 16.7% for SF decoding, which was equal to chance levels. The different features excelled with varying degrees of success, in particular in (C), spike counts marginally outperformed STMUA (p>0.05, one-sided sign test, z= −0.60, n=1200, bootstrapped, *α* = 0.05, Bonferroni corrected), but both STMUA and spike count outperformed the population rate (one-sided sign test, STMUA: z = 32.06, p = 9.3e-226 and SC: z = 31.14, p = 3.4e-213, both *α* = 0.05, Bonferroni corrected, n=1200, bootstrapped). However, for direction decoding, STMUA features performed significantly better than spike count (one-sided sign test, z = 34.06, p = 1.3e-254, n=1200, bootstrapped) and population rate (one-sided sign test, z = 34.61, p = 8.3e-263, *α* = 0.05, Bonferroni corrected, n=1200, bootstrapped), whilst spike count and population rate did not differ significantly (one-sided sign test, z = −1.99, p >0.05, n=1200, bootstrapped).

## Discussion and conclusions

We performed in-vivo extracellular electrophysiological experiments in left hemisphere V1 of the anaesthetised mouse to address questions about stimulus information at the earliest cortical processing stage. Analysis revealed that left forward drifting gratings consistently evoked the highest neuronal responses. This is a result at odds with the widespread understanding of random organisation of orientation selectivity in rodents. Spatial and directional tuning properties of electrode sites appeared very similar among animals, with high signal correlations of the tuning functions, and lower noise correlations. Further, preferred direction was affected by the SF. Finally, we showed that despite this bias in selectivity it is possible to decode stimuli from the neural responses. Here, direction decoding of spatiotemporal features attained much higher classification rates than population or spike count features, whereas spike count features attained better results than the other features in decoding SF. This suggests that these stimulus properties may be encoded differently.

As already described by (Ringach, Mineault, Tring, Olivas, Garcia-Junco-Clemente, and Trachtenberg 2016) who found that the direction tuning similarity decreased with cortical distance, this study presents a high tuning stimilarity between sites (both SF and direction) across different locations, that was also found to decrease with cortical distance, in contrast to studies that did not indicate clustering of orientation preference (Ohki, Chung, Ch’ng, Kara, and Reid 2005). In addition, we found an overrepresentation of preferred responses to leftward moving gratings (equivalent to an object moving from right temporal to nasal visual field) in left V1. We showed that preferred directions, and least-preferred directions (distribution of minimum response) were not uniformly distributed. In monkey V1, it was shown that orientation selectivity was uniformly distributed across layers (Ringach, Shapley, and Hawken 2002) whilst the distribution of orientation preference was not (Gur, Kagan, and Snodderly 2005). This overrepresentation does not necessarily contradict the current understanding of rodent V1 being randomly organised, as it could be seen as a salt-and-pepper organisation with particularly large “pepper” grains representing (in this case) leftward moving gratings. It has been known that cardinal directions exhibit stronger or more stereotyped responses and that V1 integrated locomotor and optic flow information (Saleem, Ayaz, Jeffery, Harris, and Carandini 2013; Dyballa, Hoseini, Dadarlat, Zucker, and Stryker 2018), yet, our data showed visual movements in the opposite direction of the optical flow during running evoke highest responses. Rightward moving objects, e.g. during forward movement (roughly corresponding to 0°), might thus be the more commonly observed visual stimulus, but maybe not the behaviourally most relevant.

Rodents are prey animals and close to the ground, and their eye position is more lateral and superior than e.g. cats, which enables them to identify and flee when pursued by a predator, which may come from the back, and most certainly from above. Looming stimuli and overhead sweeping discs were indeed shown to induce flight or freeze responses in mice (Yilmaz and Meister 2013; De Franceschi, Vivattanasarn, Saleem, and Solomon 2016) and thus their detection is very behaviourally relevant. The inward drifting gratings at a large SF resembling an object moving into the field of view are perhaps the most resemblant to such looming stimuli, and as such similar to the “attacking” direction from behind. The dip in 90° might be less behaviourally relevant, as an object or the world moving upwards are cases not frequently observed or simply do not pose a threat to the animal. This may play an important role in the detection of predators (i.e. it would be easier to detect a target stimulus that approaches from behind rather than one the mouse approaches).

The high correlation of tuning functions contradicts the widespread understanding of random connectivity in mouse V1, however recent studies, e.g. (Kondo, Yoshida, and Ohki 2016; Ringach, Mineault, Tring, Olivas, Garcia-Junco-Clemente, and Trachtenberg 2016; Jimenez, Tring, Trachtenberg, and Ringach 2018), also support that there may be more structure in V1 than previously reported. In particular, Ringach (Ringach, Mineault, Tring, Olivas, Garcia-Junco-Clemente, and Trachtenberg 2016) reported in a Ca^2+^ study that tuning similarity depended on cortical distance, which our findings support beyond layer 2/3 and over greater distances; and Jiminez (Jimenez, Tring,Trachtenberg, and Ringach 2018) suggested similar tuning was found when neurons responded to the same part of the visual field. The modes of directions that elicited maximum response appeared to smoothly transition from 180 to 360°. This suggests that if direction selectivity varies with SF, it could be possible to encode both size and direction information in the same population, making it an efficient coding strategy. Orientation and direction selectivity was also subject to the underlying SF used, a result which has recently been reported by (Ayzenshtat, Jackson, and Yuste 2016) in Ca^2+^-Imaging study of Layer 2/3 neurons. Selectivity indices estimated at 0.01 cpd resulted in much higher values than pooled across SFs, further arguing for a dependency on SF. Activity recorded from mouse V1 was selective to SFs and exhibited either bandpass or low-pass properties. The site studies could confirm the median preferred spatial frequency to lie around 0.02 cpd, consistent with data reported for single units elsewhere (Niell and Stryker 2008; Gao, DeAngelis, and Burkhalter 2010; Prusky, Alam, Beekman, and Douglas 2004; Umino, Solessio, and Barlow 2008; Vreysen, Zhang, Chino, Arckens, and Van den Bergh 2012). Peak SFs revealed a bimodal distribution, with one peak at the lowest SF, and the secondary peak at 0.03 cpd. This is in accordance with the observed bandpass (peak unequal to zero) and low-pass (monotone decreasing) properties. The median −3 dB cut-off frequency lay at 0.12 cpd, which is exactly the SF limit used in the literature to generate noise movies that drive a maximum number of neurons (Niell and Stryker 2008).

The use of MUA to classify multiple visual stimuli was demonstrated. A number of techniques could have been used to investigate this question. Spike-sorting the MUA signal to produce single-unit activity (SUA), while enhancing the interpretability of single cell properties, is less sensitive to the representation of features on the 100 *µ*m scale in which we are interested here. Similarly, two-photon imaging based methods are able to record from multiple individual cells simultaneously, albeit all from the same or similar superficial cortical depths. While volumetric 3D two photon imaging has advanced in recent years (Schultz, Copeland, Foust, Quicke, and Schuck 2017), it does not yet provide the ability to record from all layers and different cortical columns at the same time, while summing over as much cortical volume as attainable that is possible with our arrangement of MUA. We achieved 73% accuracy against chance levels of 12.5% solely using STMUA features. Median performance exceeded minimal correct classification rates to assert statistical significance (Combrisson and Jerbi 2015) in all cases and for all features. In SF decoding, spike count and STMUA features achieved similar correct prediction rates, whereas direction decoding achieved best results using STMUA. This implies that SF and direction features may be encoded differently. Given that SF is a static feature while direction itself is a vector quantity (i.e. time is required to perceive the movement), it may be sufficient for SF to be represented as a (static or integrated) spike count feature, whereas directions require temporal information to sufficiently decode the stimulus.

Together, the findings propose MUA in V1 contains substantial information about stimulus structure and that they can be used to efficiently decode direction and SF, potentially a useful tool for investigating changes in cortical circuit representation in behavioural learning paradigms. In addition, mouse cortex may in fact be more similar in their processing strategies to monkey and cat than previously postulated (Juavinett and Callaway 2015).

## Acknowledgements

This research was supported by the Biotechnology and Biological Sciences Research Council (BBSRC grant BB/K001817/1), and an Imperial College Department of Bioengineering PhD scholarship. We thank Dr. S Mitolo for drawing the mouse in Fig. 1 (A), Dr. S Ardila Jimenéz for useful discussions, and Dr. J Sollini for useful feedback on a draft of the manuscript.

